# Analysis of Nematode Ventral Nerve Cords Suggests Multiple Instances of Evolutionary Addition and Loss of Neurons

**DOI:** 10.1101/2025.03.20.644414

**Authors:** Jaeyeong Han, Alyson Ficca, Marissa Lanzatella, Kanika Leang, Matthew Barnum, Jonathan C. T. Boudreaux, Nathan E. Schroeder

## Abstract

Despite their diversity in habitats, nematodes are often considered to have a highly conserved neuroanatomy. This premise is based on only a subset of the nematode phylogenetic tree focused on the more diverged clades within the class Chromadorea, which includes the model organism *Caenorhabditis elegans,* thereby limiting our understanding of macroevolutionary trends in nervous system structure. To approach this problem, we used nuclear morphology to quantify the number of neurons in the nematode ventral nerve cord (VNC) to identify evolutionary patterns in neuroanatomical organization within the basal clades of Chromadorea and Enoplea. DAPI staining revealed significant VNC neuron count variations among families, with notable differences between the classes Enoplea and Chromadorea and among Enoplean species. These results may indicate a degree of evolutionary morphological stasis in later diverging Chromadorean clades. To further examine developmental patterns and potential variation in Enoplean nervous system architecture, we established an isogenic culture of the nematode *Mononchus aquaticus.* We found that while *M. aquaticus* contained four times as many VNC neuronal nuclei as *C. elega*ns, the VNC had a similar developmental timeline during post-embryonic stages. However, dye-filling assays revealed an extensive distribution of neurons along the lateral body wall, which have no obvious homolog to *C. elegans*. We found that *M. aquaticus* is capable of sustained movement following bisection that may imply a more distributed nervous system network. Our results provide a roadmap for understanding phylum-wide nervous system evolution and demonstrate large-scale differences between Enoplean and Chromodorean nervous systems.

## 1 Introduction

Nematodes are among the most widespread and diverse animals on Earth, occupying a vast range of ecological niches. Despite their diversity, research on the nematode nervous system is dominated by studies of *Caenorhabditis elegans*. While there is a staggering amount of information on the *C. elegans* nervous system, it seems unlikely that the *C. elegans* nervous system has been replicated exactly across the estimated 500 million plus years of nematode evolution (Qing et al. 2024). Recent years have seen an increased interest in understanding the nature and mechanisms of nervous system evolution in nematodes (Han et al. 2016; Toker et al. 2024; Wang et al. 2025). These studies have pointed to distinct evolutionary changes which set the stage for the use of nematodes to dissect broad principles of nervous system evolution.

Phylogenetic trees based on 18s ribosomal DNA and genome sequences support the division of the phylum Nematoda into the classes Enoplea and Chromadorea (Figure 1; Qing et al. 2024). These have been further subdivided into twelve clades (Holterman et al. 2006; van Megen et al. 2009). From these data, a consensus has emerged that the subclass Enoplia (Clade 1) is likely the earliest diverging nematode clade, followed by Dorylaimia (Clade 2). Nematodes in these earlier diverging clades differ from Chromadorean clades in several aspects of their morphology and development (Schulze and Schierenberg 2011). For example, several species in the basal clades lack the deterministic development seen in embryonic development of *C. elegans* (Clade 9) and other species in more recently derived clades (Schulze and Schierenberg 2011).

**Figure 1.**
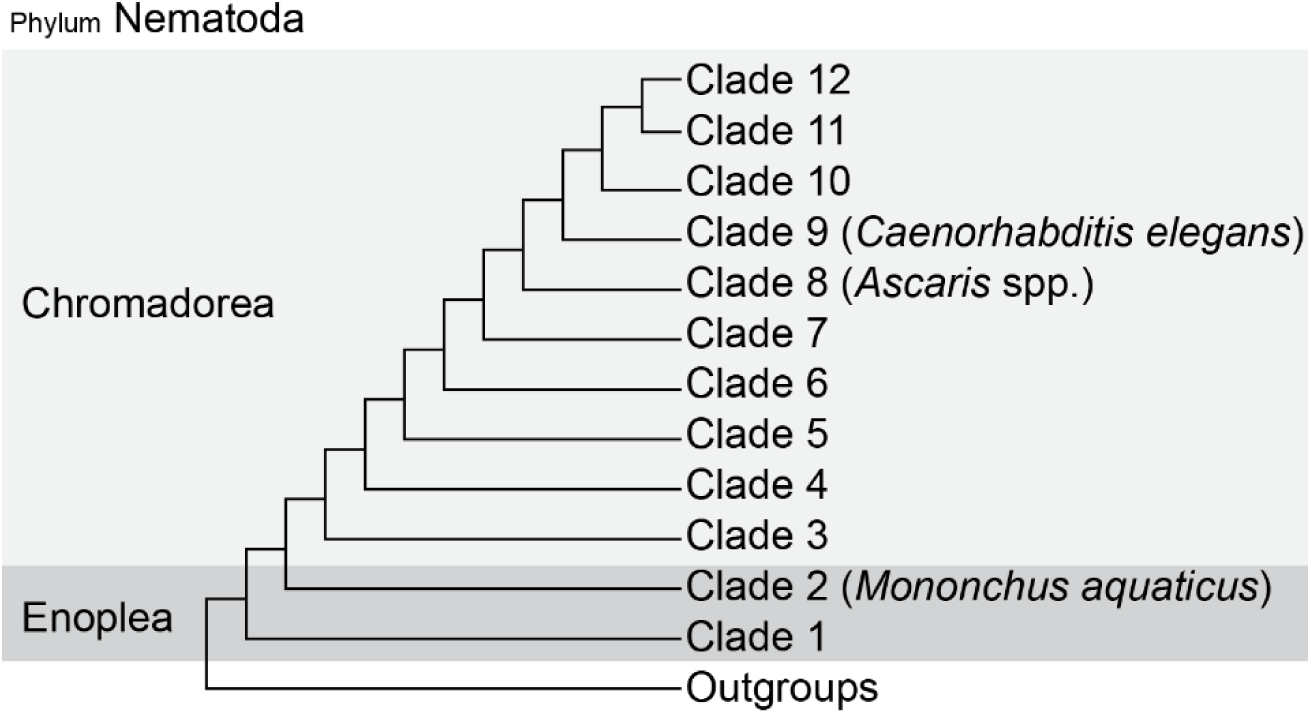
Modern cladistic phylogeny of nematode adapted from Holterman et al. (2006) and van Megen et al. (2009). Note that branch lengths do not represent evolutionary distance.

The remarkable similarity in neuroanatomy between *C. elegans* (Clade 9) and the gastrointestinal parasite *Ascaris suum* (Clade 8) suggests a high degree of nervous system conservation. Despite distinct habitats, sizes, and an estimated evolutionary separation of 350 million years, these species have nearly identical numbers (∼300) of neurons (Qing et al., 2024; Stretton et al. 1978; Sulston 1976; White 1976). Recent electron microscopy data comparing *C. elegans* to the satellite species *Pristionchus pacificus* (Clade 9) also found a nearly identical number of neurons in the anterior nervous system (Cook et al. 2025). However, these more detailed EM results also demonstrated instances of large-scale rewiring. Previous light-microscopy from our work compared nine nematode species within clades 9-12 and found differences in the number of putative neurons within the ventral nerve cord (VNC) suggesting a previously unappreciated amount of neuroanatomy diversity among a subset of the phylum (Han et al. 2016). One limitation from these studies is that they exclude a large portion of the phylum Nematoda from clades 1-6 (herein referred to as “basal clades”).

Due to their relative recalcitrance to culturing, very few studies have examined the neuroanatomy of nematodes in basal clades. The handful of investigations suggested that certain basal clade nematodes may have an order of magnitude more neurons than found in *C. elegans* and other later diverging clades (Gans and Burr 1994; Malakhov 1994; Sulston and Horvitz 1977). These findings could suggest a possible secondary simplification where the number of neurons decreased at some point during the evolution of the Chromadorea. To test the hypothesis of nervous system simplification between Enople and Chromadorea, we used a survey approach to quantify the number of neurons in the VNC across clades 1-6, representing an estimated 500 million years of evolution (Qing et al. 2024). As this survey approach prevented an examination of intraspecies variability, we also established an isogenic culture of *Mononchus aquaticus* (Clade 2). By comparing our findings with existing knowledge on Chromadorean species such as *C. elegans*, we highlight both conserved and divergent neuroanatomy across the phylum.

## 2 Results

### 2.1 Total Number and Density of VNC Neuron-like Nuclei Does Not Support a One-time Simplification Process

Similar to our previous work (Han et al. 2016), we used nuclear morphology as an indicator of cell type in order to enumerate putative neurons in the VNC of multiple nematode species. However, as few species in basal clades are considered culturable, we performed DAPI (4’,6-diamidino-2-phenylindole) staining on nematodes directly recovered from diverse habitats (Table 1). Individuals were identified to the family level and the number of VNC neuron-like nuclei were counted between the retrovesicular ganglion (RVG) and preanal ganglion (PAG) (Figure 2). VNC neuron data were compared across multiple taxonomic levels, including family, clade, and class. While there was a significant difference between Enoplean and Chromadorean nematodes (Figure 3a; P<0.0001), we noted substantial variability within each of these groups. At the clade level, VNC neuronal nuclei counts differed significantly (P<0.0001; Figure 3b). Clades 2 (Dorylaimia) and 6 exhibited significantly higher number of neurons than other clades, with clade 2 showing significantly higher numbers compared with clades 3, 5, 9, 10, 11, and 12. The large number of neuronal nuclei in clade 6 was solely due to individuals from Camacolaimidae isolated from marine samples. At the family level, we also observed significant variation in VNC neuronal nuclei counts (P < 0.0001,Figure 3c).

**Figure 2.**
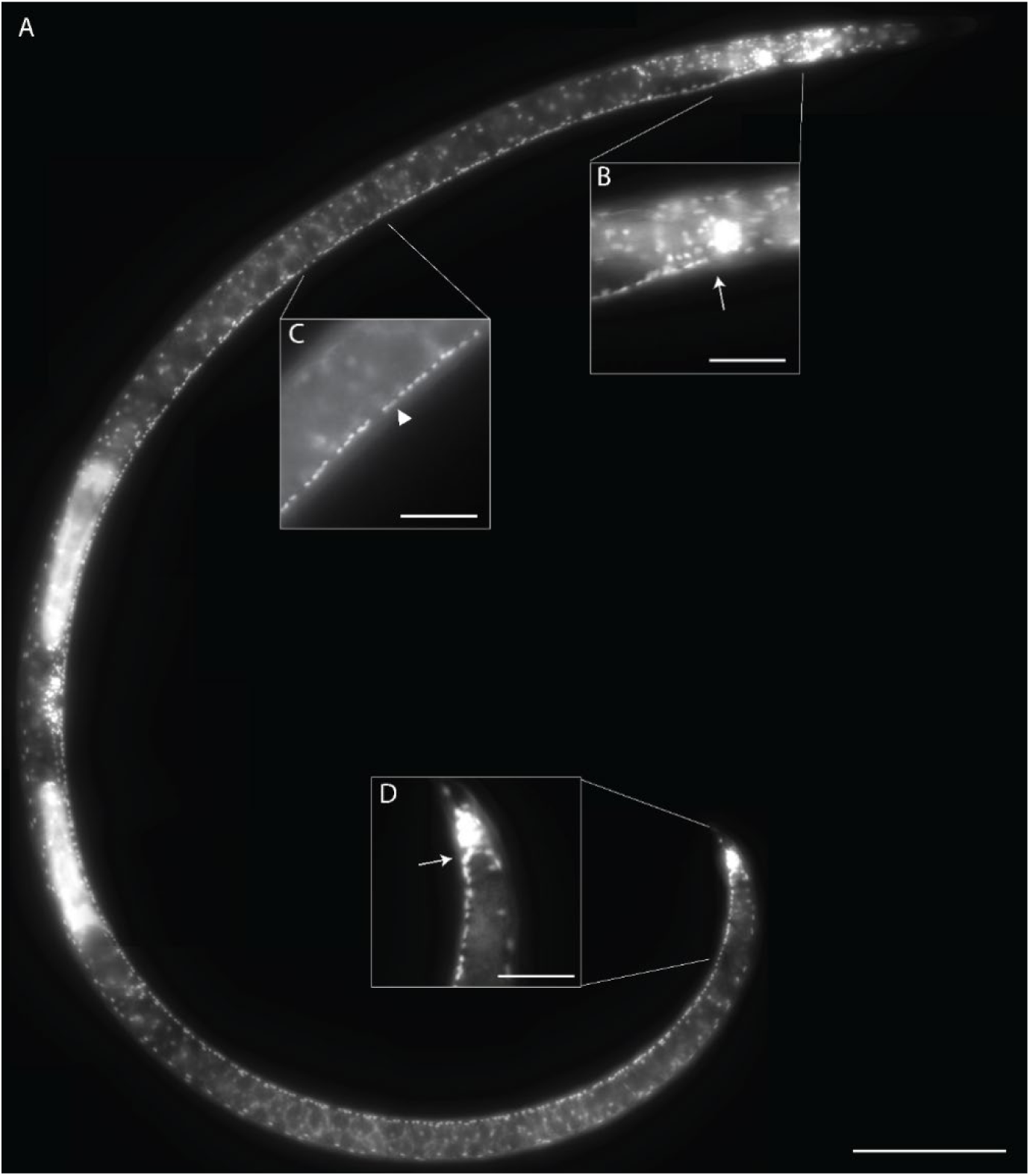
DAPI staining of *Xiphinema* sp. (Clade 2). The ventral nerve cord (VNC) consists of a line of neurons extending along the ventral line from the retrovesicular ganglion (RVG) to the pre-anal ganglion (PAG). (A) Lateral view of an adult female *Xiphenema* showing overall VNC structure. (B) Close-up of the region at RVG and the anterior VNC (arrow). (C) Putative hypodermal (arrowhead) and adjacent neuronal nuclei of VNC. (D) Division between PAG and VNC (arrow). Scale bars = (a) 100 µm; (b-c) 25 µm.

**Figure 3.**
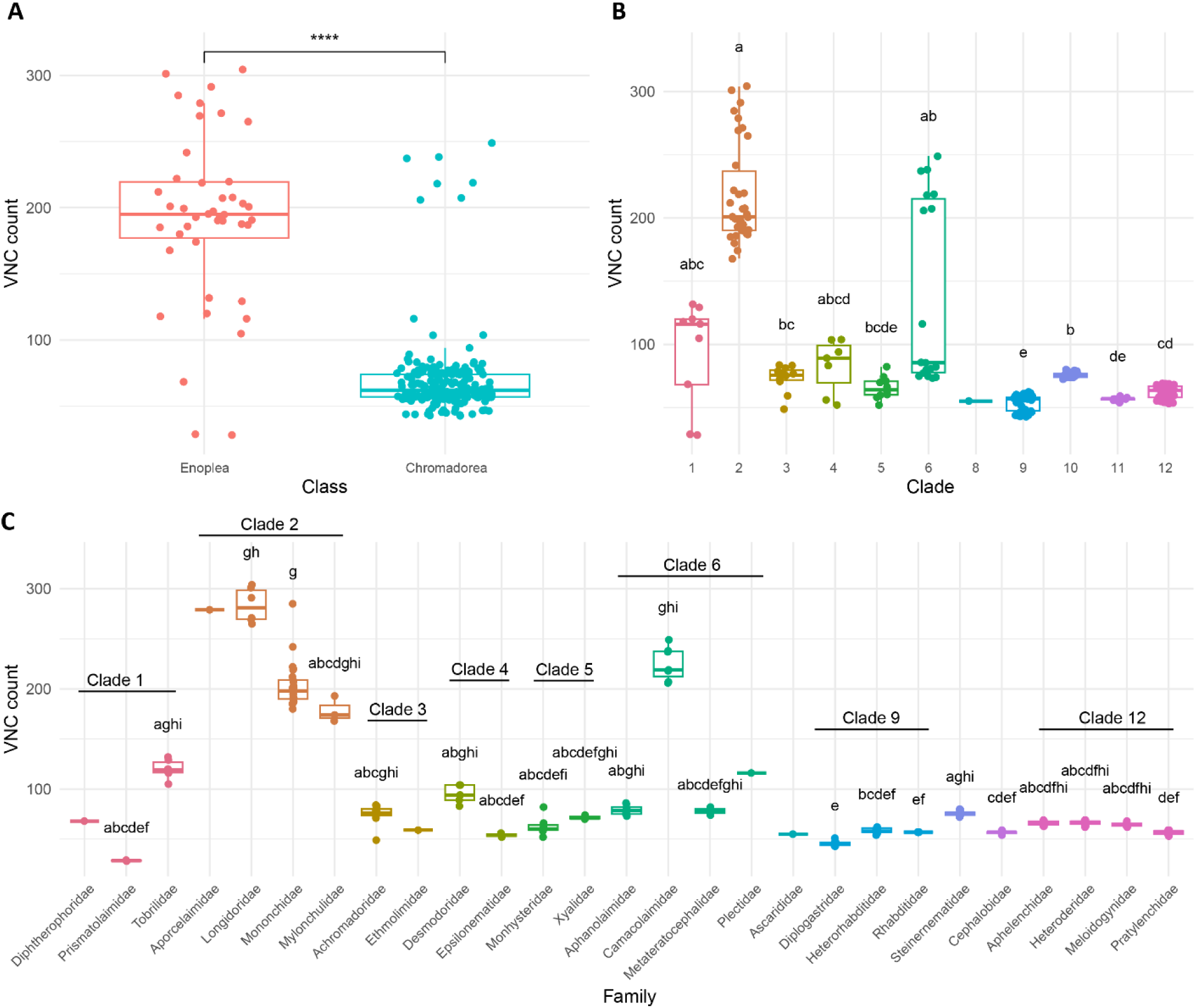
Analysis of VNC neuronal nuclei counts across family (A), clade (B), and class (C) levels. Clades 1-6 include single female adults except for Camacolaimidae and Epsilonematidae, which included males. Clades 9-12 data, previously published in Han et al. (2016), included several taxa where juveniles were examined. Families with a sample size of 1 (Aporcelaimidae, Ascarididae, Diphtherophoridae, Ethmolaimidae, and Plectidae) were excluded from family level comparison. Groups sharing the same letter designation are not significantly different according to Dunn’s multiple comparisons test at α=0.05 (A and B). A Student’s t-test was performed for class-level comparison (C). **** indicates P ≤ 0.0001.

Despite having hugely different body sizes and numbers of muscle cells, *C. elegans* and *A. suum* have nearly identical numbers of VNC motor neurons (Stretton et al. 1978), suggesting that number of neurons does not necessarily need to scale with body size. However, we wanted to determine if there was a relationship between body size and VNC neurons outside of the Chromodorea. Indeed, regression analysis suggested a moderately strong positive correlation between VNC length and number of neuronal nuclei (R^2^ = 0.64, Figure 4). We found a statistical difference in VNC neuron density between classes with Enoplea being higher than Chromadorea (P<0.0001; Figure 5.a). However, similar to examination of total number of neuron-like nuclei, we also found a statistically signficant difference in VNC neuronal density across all clades and between select pairwise family comparisons (Figure 5.b and c). At the clade level, differences in neuron density were less pronounced than those observed with total VNC neuron counts (Figure 5b). For example, while clade 2 had a higher density than clades 1, 4, and 9, it was not significantly different from the other clades unlike the results from total neuron counts (Figure 3b). At the family level, Longidoridae had the highest neuronal density (neuronal nuclei per 100 µm), significantly higher than Achromadoridae, Aphanolaimidae, and Rhabditidae (Figure 5c). Rhabditidae (*C. elegans* only) had the lowest density and significantly lower, compared with Longidoridae, Mononchidae, Mylonchulidae, Monhysteridae, Camacolaimidae, and Metateratocephalidae. Together, these results do not support a one-time simplification of neuroanatomy, but point to more complex trends in the evolution of nematode nervous systems with both streamlining and expansion of neuronal numbers.

**Figure 4.**
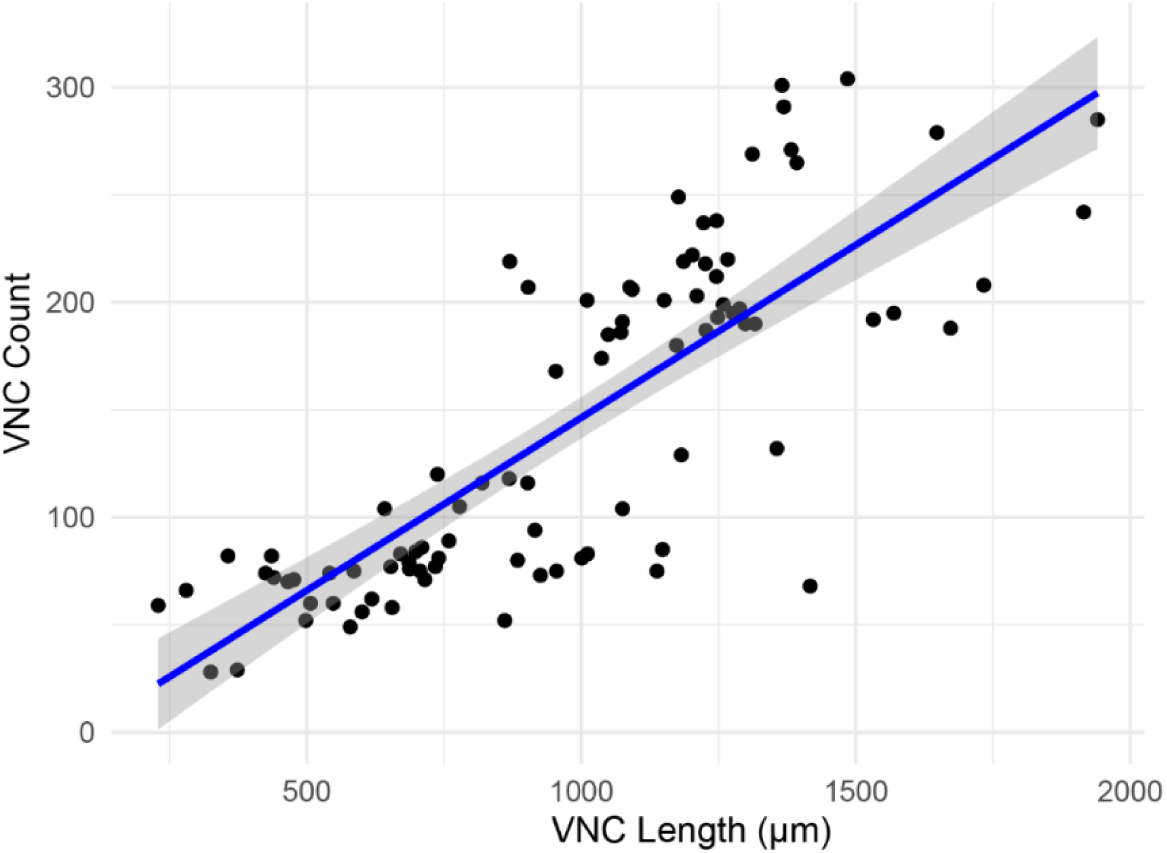
Linear regression analysis of VNC neuronal nuclei counts on VNC length (µm). Each point represents a measurement of a single female from Clades 1-6 except for Camacolaimidae and Epsilonematidae, which included males. The regression line shows a positive correlation between VNC length and neuronal nuclei count (slope: 0.1607, adjusted R^2^: 0.64, P < 0.001).

**Figure 5.**
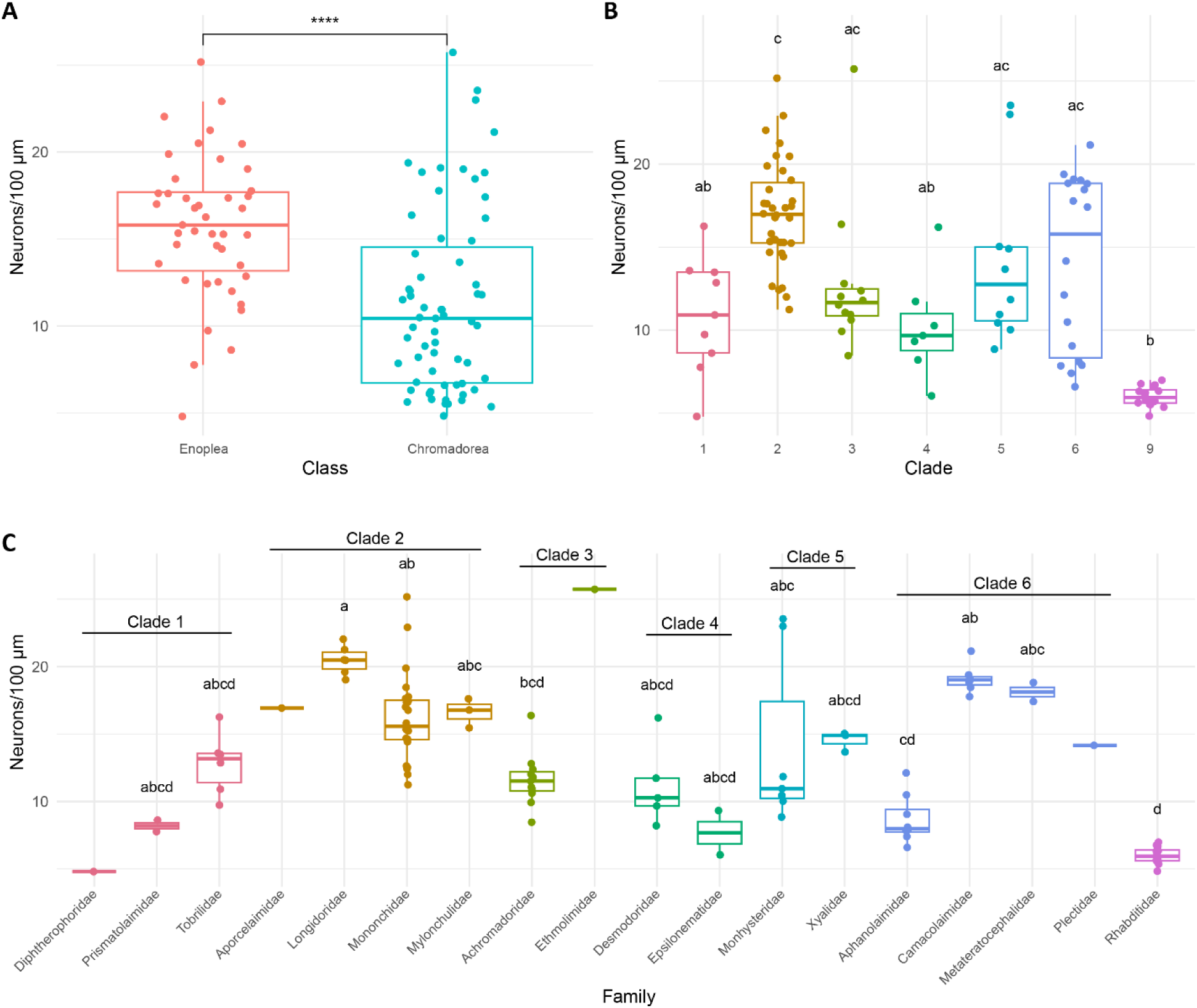
Analysis of VNC neuronal nuclei density (neurons per 100um) across family (A), clade (B), and family (C) levels. Clades 1-6 include single female adults except for Camacolaimidae and Epsilonematidae, which included males. Clades 9-12 data, previously published in Han et al. (2016), included several taxa where juveniles were examined. These were, therefore, excluded from density analyses except *Caenorhabditis elegans* (Rhabditidae, Clade 9) where sufficient females samples were examined for analysis. Families with a sample size of 1 (Aporcelaimidae, Ascarididae, Diphtherophoridae, Ethmolaimidae, and Plectidae) were excluded from family-level comparison. Groups sharing the same letter designation are not significantly different according to Dunn’s multiple comparisons test at α=0.05 (A and B). A Student’s t-test was performed for class-level comparison (C). **** indicates P ≤ 0.0001.

### 2.2 *Mononchus aquaticus* Exhibits Determinate-Like VNC Development

Among the various survey data, we noted that the Dorylaimia (clade 2) had more VNC neuronal nuclei than other taxa in our survey. Within this subclass, previous studies have reported on the successful cultivation of Mononchida nematodes (Grootaert and Maertens 1976; Maertens 1975; Nelmes 1974; Salinas 2004). Mononchida comprises predatory nematodes characterized by a large dorsal tooth. To better examine nervous system development and variability among individuals, we established an isogenic culture of *Mononchus aquaticus* from a single female. We counted 198±12 (n=20) neuronal nuclei in the VNC of adult *M. aquaticus* females (Figure 6).

**Figure 6.**
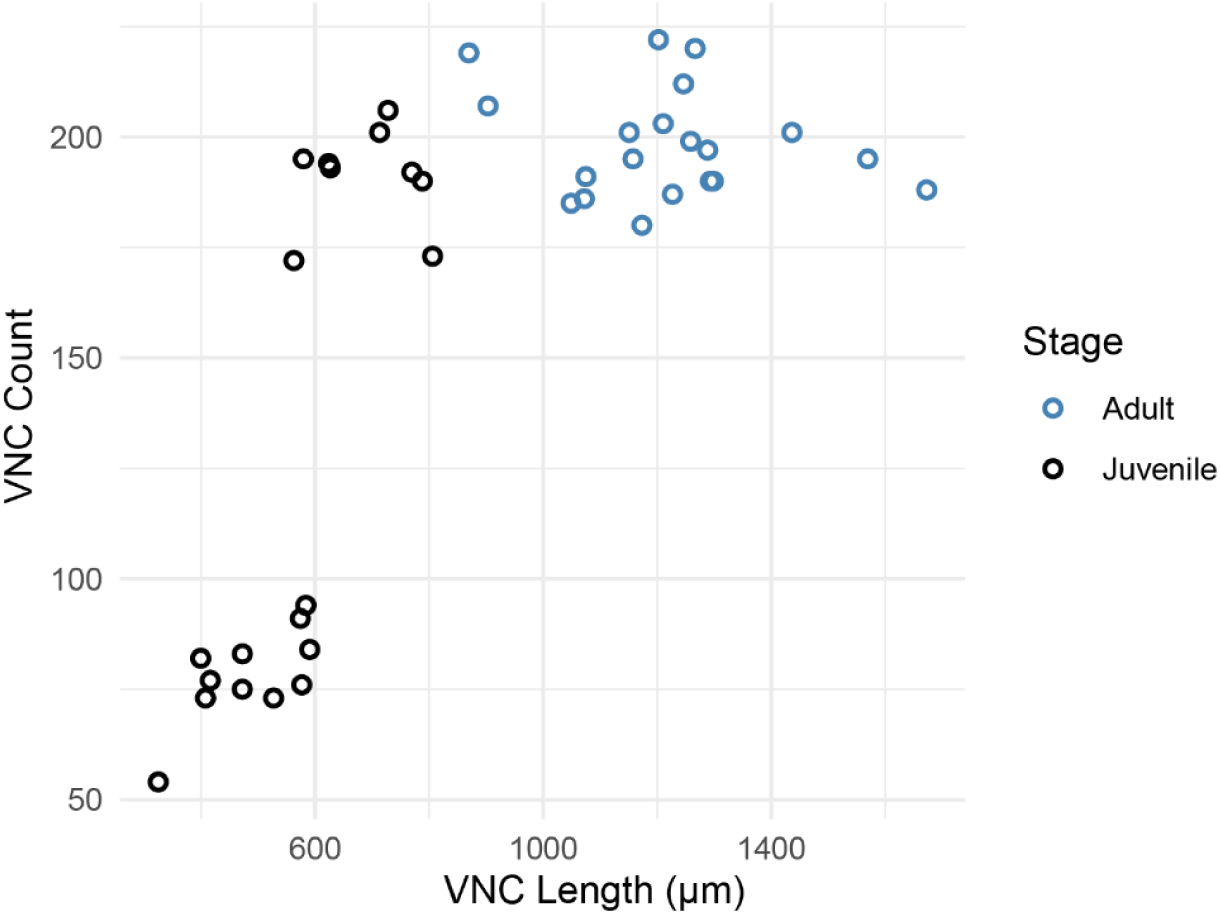
VNC neuronal nuclei counts through the development of cultured *Mononchus aquaticus*. A positive association between VNC neuronal nuclei counts and VNC length was found in juveniles (P<0.0001, adjusted R^2^: 0.607, slope: 0.33825), while no association was found in adults (P=0.03595, adjusted R^2^: −0.00613, slope: −0.01385). Black circles represent juveniles; blue circles represent adults.

In *C. elegans,* only a subset of VNC neurons form during embryogenesis. This is followed by the rapid addition of additional neurons and rewiring during late L1 (Sulston 1976). Similar developmental timelines were found in several other Chromodorean species (Bui and Schroeder 2018; Han et al. 2016). We wished to see if a similar process occurs in an Enoplean species. As we have not strictly determined the developmental timeline for *M. aquaticus,* we used VNC length as a surrogate for development. Our results suggest a similar pattern of VNC development as found in *C. elegans* (Sulston and Horvitz 1977). The smallest juveniles (presumed J1s) had counts of 54-94 VNC neuron-like nuclei (Figure 6). However, this number increased rapidly during juvenile development to nearly 200 VNC neuron-like nuclei. No correlation between VNC length and number of neuron-like nuclei was found following this rapid increase.

### 2.3 Dye-Filling Reveals Expanded Neuron Distribution in *M. aquaticus*

As nuclear morphology may not be accurate in assessing cell type, we also deployed a dye-filling assay commonly used to assess sensory neurons in *C. elegans* (Tong and Bürglin 2010). In *C*. *elegans*, six pairs of amphid head neurons and two pairs of phasmid tail neurons, including cell bodies and neuronal processes fluoresce following exposure to specific lipophilic dyes (Tong and Bürglin 2010). While some variation exists, dye filling among Chromodorean nematodes is always restricted to neurons in the anterior and posterior ends (Garg et al. 2022; Han et al. 2016; Srinivasan et al. 2008).

In addition to dye-filling in anterior and posterior neurons, we also observed large numbers of dye-filled neurons along the entire length of the *M. aquaticus* lateral body walls (Figure 7). The anterior neurons were similar to the position and morphology of both the amphid and inner-labial neurons of *C. elegans.* Staining of the circumpharyngeal nerve ring appeared as two distinct anterior and posterior regions. The specific morphology of the body wall neurons varied. All body wall neurons included at least two processes emerging from the cell body; however, many neurons included three or four processes. In all instances, at least one process traveled to the VNC. Occasionally, both processes of bipolar neurons entered the VNC. We also observed occasional commissures where neuronal processes traveled from the dorsal to ventral cords without an apparent cell body (Figure 7d). As we never observed dye-filled cell bodies within the VNC or dorsal cord (DC), these commissures likely originated from a cell body in the lateral body wall that sent a process into the VNC or DC, which then extended back to the opposite cord. While most body wall neurons within the lateral body wall were separate enough to discern individual cell bodies and processes, we also regularly observed clusters of four to five body wall neurons in close enough proximity that we could not determine the source of individual processes. In total, we counted 120±36 (n=16) dye-filled neurons in *M. aquaticus* adults (Figure 9a) in contrast to the 16 neurons that typically dye fill in *C. elegans*.

**Figure 7.**
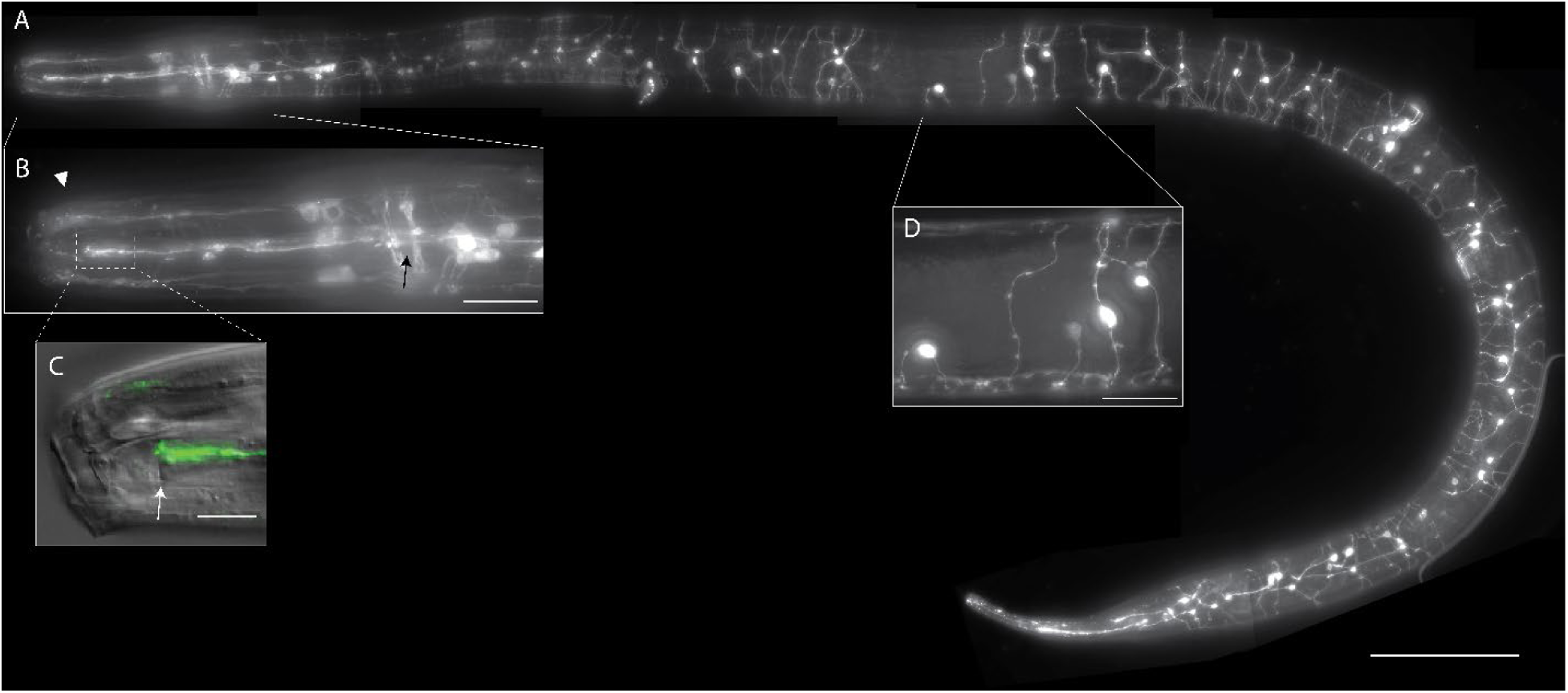
Dye-filling assay in adult *Mononchus aquaticus.* (A) Lateral view of entire body of dye-filled female *M. aquaticus*. (B) Dye-filled head sensory neuron cell bodies and the nerve ring (arrow). Putative inner labial and amphid neuron dendrites are extended to the anterior end. The nerve ring appears to be separated into two bundles and receives processes from surrounding neurons. (C) Amphid neuron dendrites terminate at the slit-shaped amphid aperture (arrow). (D) Body wall neurons distribute throughout the body, with processes extending to the ventral and dorsal nerve cords to form commissures. Scale bars = (a) 100 µm; (b, d) 25 µm; (c) 10 µm.

The nervous system of *C. elegans* is highly stereotyped. While many *C. elegans* neuron cell bodies show some positional variability, this is typically on the order of 10 µm (Toyoshima et al. 2020; Yemini et al. 2021). The large amount of variation in dye-filling counts we observed in *M. aquaticus* could be due to both variation in uptake of dye by individual neurons and differences in the number of neurons between indviduals. To determine the consistency of staining we compared cell body position, number of processes and process directionality of body wall neurons within a defined 50 µm area in multiple individuals. As a landmark we selected the 50 µm surrounding the vulva. Only animals with bright staining in other parts of the nervous system were selected. We found that each animal had a distinct number of neurons and branching pattern suggesting a high degree of inter-individual variability in body wall neuroanatomy (Figure 8).

**Figure 8.**
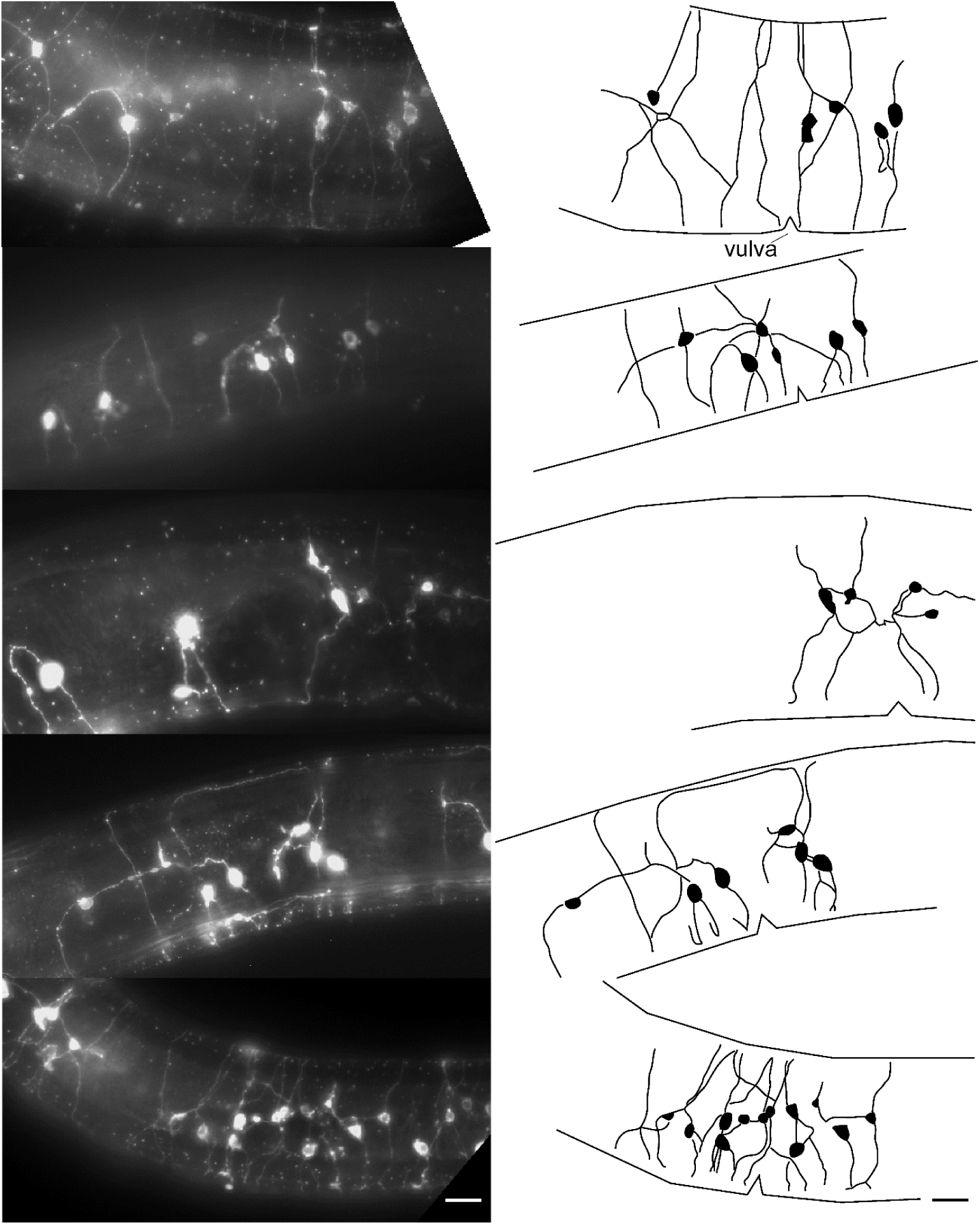
Lateral view images of dye-filled near the vulval region in adult *Mononchus aquaticus* (Left). GFP fluorescence shows the arrangement of neuronal cell bodies and processes. All nematodes are displayed with anterior oriented to the left and the dorsal side up. (Right) Corresponding schematic outlines of neurons (black fill) and processes adjacent to the vulva. Processes are not drawn within the ventral or dorsal cords as it was impossible to differentiate individual neurons within the cords. Scale bar = 10 µm.

Similar to VNC neuron counts, we found more dye-filled neurons in adults compared to juveniles (Figure 9a). Interestingly, we found that the head sensory neuron numbers remained consistent throughout development (Figure 9b), suggesting that head sensory neurons may form during embryogenesis while body sensory neurons are continuously added during post-embryonic development as the nematode increases in length. When examined as a function of total body length, juvenile dye-filled neuron numbers positively correlated with VNC length (R^2^ = 0.52) but plateaued in adults (R^2^ = 0.13) (Figure 9c). While similar to the developmental timeline of VNC neuronal nuclei (Figure 6), dye-filled neurons showed gradual addition during juvenile development rather than a sudden increase.

**Figure 9.**
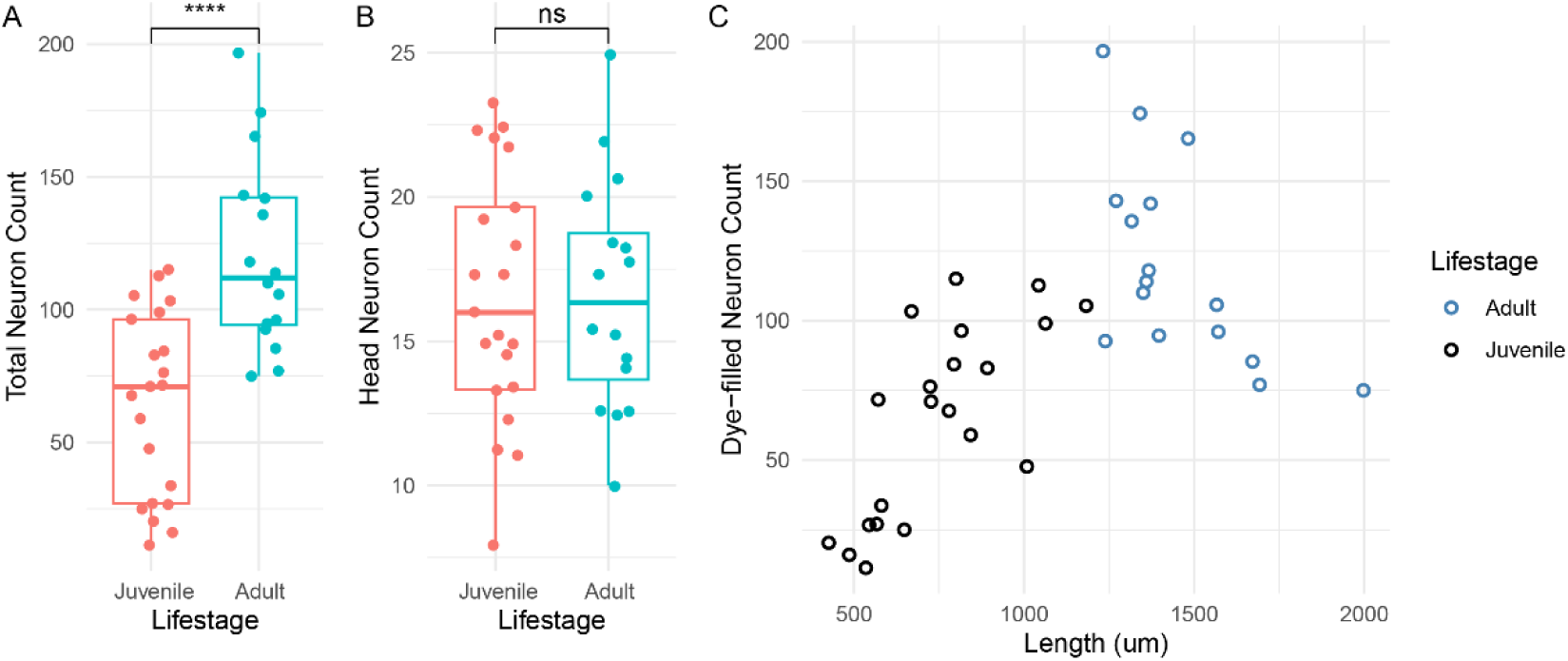
Comparative analysis of dye-filled neurons in *Mononchus aquaticus* across developmental stages. (A) Total dye-filled neuron counts in adults versus juveniles. Adults exhibit significantly higher total neuron counts (120±36, n=16) compared to juveniles (64±35, n=21; P < 0.0001, t-test). (B) Dye-filled head neuron counts in adults (17±4, n=16) versus juveniles (16±4, n=21). No significant difference was detected (P=0.9966, t-test). (C) Relationship between total dye-filled neuron count and body length (µm) in *M. aquaticus*. While there is a moderately positive association of dye-filled neuron counts and body length in juvenile (P < 0.0001, adjusted R^2^: 0.4984, slope: 0.12156), weak negative association was found in adult (P< 0.05, adjusted R^2^: 0.3299, slope: −0.10817).

### 2.4 Decentralized post-bisection response in *M. aquaticus*

Our results here suggest that *M. aquaticus,* as well as other Enoplean nematodes, have substantially more neurons than *C. elegans.* An obvious question is whether there are any functional outcomes for these additional neurons? Anecdotally, we did not observe any obvious behavior novelties in *M. aquaticus* compared to other nematodes; however, we made an accidental behavioral finding following bisection of animals. During the molecular identification of *M. aquaticus* we cut the animals to facilitate the cell lysis step prior to PCR. We noted that that the two halves of *M. aquaticus* kept moving rapidly even several minutes after being bisected. To test this further, we compared the body bends in the posterior halves of *M. aquaticus* and *C. elegans* before and after bisection (Figure 10). Uncut *C. elegans* and *M. aquaticus* showed average body bends per minute of 62 and 40.8, respectively. Following bisection, *C. elegans* showed an immediate reduction to 8.2 bends per minute, with near-complete cessation by 10 minutes post-bisection. In contrast, *M. aquaticus* maintained reduced but persistent body bending, showing gradual decline over 30 minutes post-bisection. These results may suggest a more decentralized nervous system in *M. aquaticus*.

**Figure 10.**
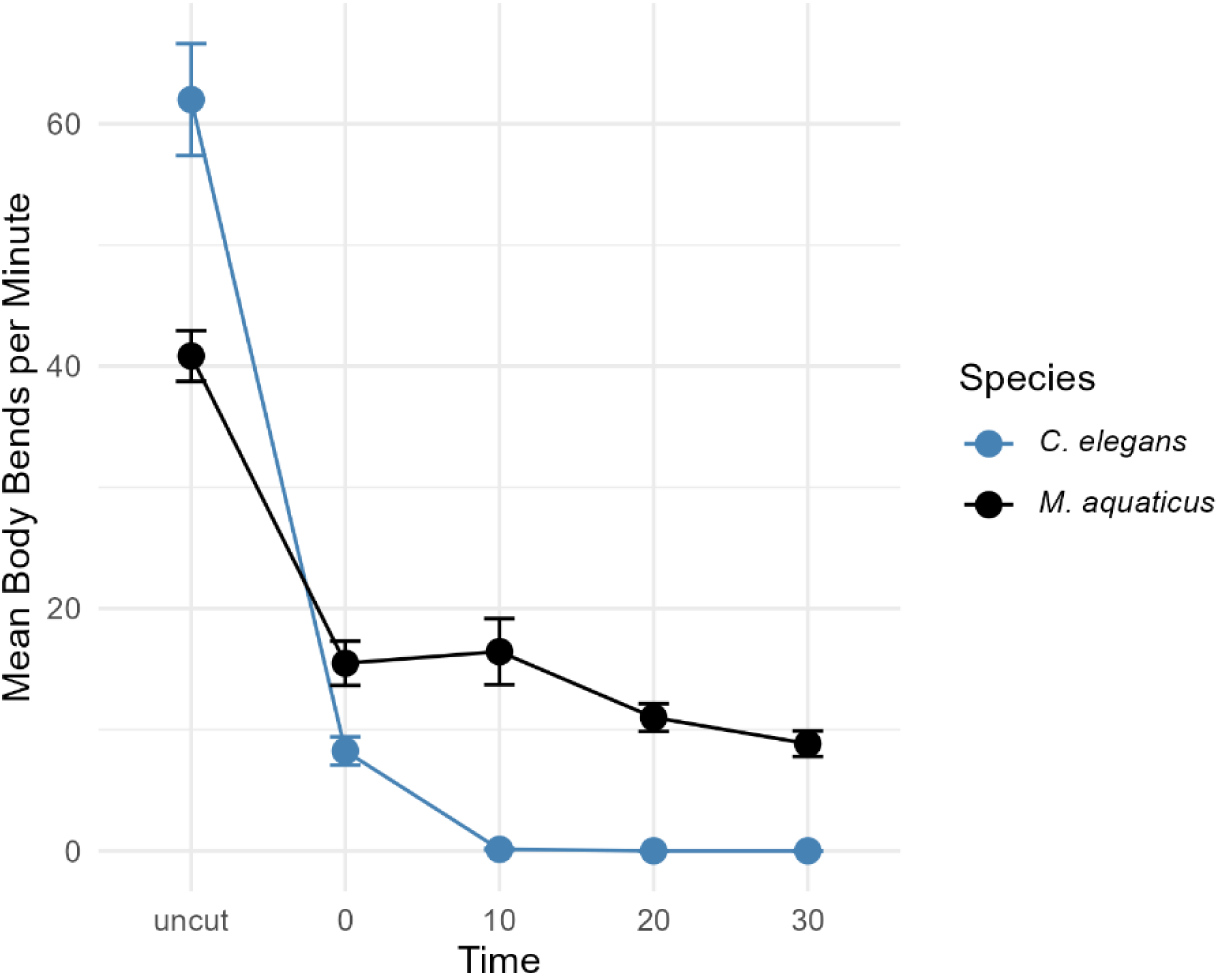
Temporal profile of body bends per minute following bisection in *Caenorhabditis elegans* (blue) and *Mononchus aquaticus* (black). Measurements were taken for uncut animals and subsequent time points (0, 10, 20, 30 minutes post bisection). Error bars represent the standard error of the mean.

## 3 Materials and Methods

### 3.1 Sample Collection and Nematode Extraction

Shallow aquatic and intermediate terrestrial sediment samples were collected and stored at 4°C until further processing. For aquatic samples, a liquid layer was always included to simulate the natural environment until processing. The majority of samples were collected from Champaign-Urbana and surrounding areas in Illinois, with additional sites in Monmouth and Belleville, Illinois (Table 1). Marine samples were collected from the Carpinteria State Beach and Salt Marsh in California, courtesy of Dr. Tiago José Pereira and Dr. Holly Bik, formerly of the University of California, Riverside.

**Table 1.** List of sampling locations for the nematodes used for DAPI staining.

| Sample Location | Locality | GPS Coordinates | Biome |
| --- | --- | --- | --- |
| Dana Colbert Park Pond | Savoy, IL | 40°03'02.1"N; 88°14'58.9"W | Freshwater |
| Prairie Meadows Subdivision Pond | Savoy, IL | 40°02'58.5"N; 88°14'47.4"W | Freshwater |
| Lake Park Residential Pond | Savoy, IL | 40°03'53.2"N; 88°14'21.1"W | Freshwater |
| Moorman Sewage Retention Pond, Large (Lake Nalbandov) | Urbana, IL | 40°05'22.2"N; 88°14'08.4"W | Freshwater |
| Moorman Sewage Retention Pond, Small (Lake Nalbandov) | Urbana, IL | 40°05'16.5"N; 88°14'11.3"W | Freshwater |
| UIUC Japan House Main Pond | Urbana, IL | 40°05'34.0"N; 88°12'59.2"W | Freshwater |
| UIUC Japan House Minor Pond | Urbana, IL | 40°05'37.4"N; 88°12'55.8"W | Freshwater |
| Boneyard Creek | Champaign, IL | 40°06'40.4"N; 88°13'40.4"W | Freshwater |
| Private Residence Pond | Urbana, IL | 40°05'49.4"N; 88°12'12.4"W | Freshwater |
| Salt Fork River, Mulberry Tree | Homer, IL | 40°05'39.1"N; 87°99'65.4"W | Terrestrial |
| Homer Lake Forest Preserve Collins Pond | Homer, IL | 40°05'39.1"N; 87°99'65.4"W | Freshwater |
| Homer Lake Forest Preserve, White Oak | Homer, IL | 40°05'39.1"N; 87°99'65.4"W | Terrestrial |
| Indian Grass (*Sorghastrum nutans*)<br>Prairie | Homer, IL | 40°05'39.1"N; 87°99'65.4"W | Terrestrial |
| Northwestern Illinois Agricultural<br>Research and Demonstration Center | Monmouth, IL | 40°93'64.6"N; 90°72'06.5"W | Terrestrial |
| Agricultural Field | Belleville, IL | 38°31'59.5"N; 89°53'40.2"W | Terrestrial |
| Carpinteria Salt Marsh, Muddy Site | El Estero, CA | 34°24'01.7"N; 119°32'21.6"W | Marine |
| Carpinteria Salt Marsh, Sandy Site | El Estero, CA | 34°23'55.6"N; 119°32'19.3"W | Marine |
| Carpinteria State Beach, Intertidal<br>Site 1 | Carpinteria, CA | 34°23'35.4"N; 119°31'30.4"W | Marine |
| Carpinteria State Beach, Intertidal<br>Site 2 | Carpinteria, CA | 34°23'35.4"N; 119°31'30.4"W | Marine |

Nematode extraction was accomplished through sieving and sugar centrifugation followed by the Baermann funnel method (MacGuidwin and Bender 2010). Nematodes were separated from sediment using 250 and 38-μm pore size sieves. Materials collected on a 38-μm pore sieve was centrifuged at 3000 rpm for 3 minutes followed by sugar centrifugation at 3000 rpm for 3 minutes with 45% sucrose. The supernatant was applied to a 25-μm pore size sieve for rinsing and collection. To prevent osmotic imbalance for marine nematodes, rinses were carried out with Instant Ocean^®^ Sea Salt, prepared as per manufacturer’s instructions.

To capture nematodes that might be missed by sieve extraction (e.g., those within roots or larger specimens), sediment collected on the 250-μm sieve was processed using a Baermann funnel apparatus (MacGuidwin and Bender 2010). After 48 hours, the sample was drained for collection. As with the sieve extraction steps, marine samples were steeped in Instant Ocean^®^ mixture to ensure cellular preservation.

Nematodes were identified to the family level based on morphological keys (Abebe et al. 2006; Bongers 1988; Mai and Mullin 1996; Peña-Santiago 2014; Schmidt-Rhaesa and Holger Rothe 2014).

### 3.2 DAPI Staining and Microscopy

Nematodes samples were centrifuged at 4000 rpm for 4 minutes in 15 ml Falcon tubes. The liquid was gently pipetted off and reduced to 1 ml. Nematodes isolated from aquatic environments tended to coil during the fixation step and, therefore, 50 μl of 0.1M levamisole was added as a relaxant. Once adequately immobilized, all liquid except a thin layer was pipetted off and 1 ml of DESS (dimethyl sulfoxide, disodium EDTA, and saturated NaCl) was added (Yoder et al. 2006). For terrestrial samples, no sedative was added and all liquid except a thin layer was immediately pipetted off and 1 mL of saturated DESS was added and stored at 4°C. Depending on the amount of sediment in each sample, 2-8 μg/mL DAPI was added to the fixed specimen. The sample was kept in the dark at RT overnight and then returned to 4°C until imaging.

DAPI-stained nematodes were mounted onto a 2% agar pad. Fluorescent microscopy was performed using a Zeiss® Axio Imager M2 fluorescent microscope equipped with an Axiocam 506 mono camera and an X-Cite® Series 120Q fluorescent lamp. Samples were screened for adults with consistent staining. Z-stacks were taken with slices at 0.35-0.5 μm intervals, collecting both the differential interference contrast (DIC) and DAPI channels for future reference. The entire nematode was imaged with sufficient overlap for subsequent stitching.

### 3.3 Ventral Nerve Cord Enumeration

Images were stitched together using linear blending and 5 peaks in Fiji (Schindelin et al. 2012). For images that were more difficult to stitch, the settings were changed to up to 50 peaks, subpixel accuracy, and zero values were ignored. The entire nematode was stitched from head to tail.

Using the stitched DAPI image, the VNC neuron-like nuclei were counted. Nuclei were considered VNC neurons based on morphology, size, position, and relative intensity of stain, in accordance with previous work (Han et al. 2016; Sulston 1976; Sulston and Horvitz 1977; White 1976). VNC neuron-like nuclei tend to be small, punctate, aligned, and brighter than surrounding nuclei (Figure 2b). Counts began at the PAG immediately anterior to the anus and proceeded towards the RVG using the multipoint tool in Fiji (Han et al. 2016). VNC length was measured from the nerve ring to the PAG, and density was calculated as neuron-like nuclei per 100 μm. To account for the possible confounding effects of sex and developmental stage, we only examined adult females with exceptions for Camacolaimidae and Epsilonematidae. For Camacolaimidae, only males were found in our sampling, while for Epsilonematidae, males were included due to low sample numbers of females and consistency between sexes.

### 3.4 *Mononchus aquaticus* Culture

*M. aquaticus* was isolated from a maize field at the Northwestern Illinois Agricultural Research and Demonstration Center in Monmouth, Illinois and identified using morphological (Ahmad and Jairajpuri 2010) and 18S sequencing results (Holterman et al. 2006; GenBank Accession: PV282461). Our isogenic *M. aquaticus* culture was established from a single female cultured based on techniques described by Salinas (2004). Briefly, soil extract agar was made by adding water to 100 cm^3^ of soil up to a total volume of 1 L and incubating overnight at room temperature. The following day, the mixture was poured through a 25-µm filter followed by centrifugation of the flow through solution for 3 minutes at 3000 rpm. The pellet was discarded and 500 mL of the supernatant was mixed with 5 g of agar and 500 µl of 5 mg/ml cholesterol dissolved in ethanol. Mixed stages of *Aphelenchus avenae,* originally isolated by Nathan Schroeder from Somerset, NJ and grown on half-strength potato dextrose agar with *Monilinia fructicola* or *Botrytis cinerea*, were used as prey. *M. aquaticus* was fed *A. avenae* weekly or biweekly. No attempt was made to ensure the absence of microbial contamination. While *M. aquaticus* readily fed on *A. avenae,* we also observed juvenile *M. aquaticus* displaying pharyngeal pumping in the absence of a prey nematode, similar to the observations made by Bilgrami (1984). *M. aquaticus* is considered parthenogenic and we have only observed a single male once.

### 3.6 Dye-Filling Assay

A total of 51 *M. aquaticus* (32 juveniles and 19 adults) were subjected to a dye-filling assay. Groups of five worms were placed in a well containing 1 mL of 5% M9 buffer. 2 µL of 2 mg/mL DiO stock solution was added to the well, and worms were incubated for 5 hours for adults or 4 hours for juveniles. After incubation, worms were transferred to empty soil extract agar plates for a 30-minute recovery period. For imaging, anesthetization was achieved by applying 10 µL of 10 mM sodium azide solution onto a 5% agar pad, onto which a single worm was placed. Images were captured using DIC and GFP filters on a Zeiss Axio Imager M2 microscope at 63× magnification.

Image analysis was performed using Fiji (Schindelin et al. 2012). Neurons were counted in the head (from the tip of the head to immediately posterior to the nerve ring) and the body (from posterior to the nerve ring to the anus). Counts were conducted using the Cell Counter plugin in Fiji. The entire nematode length was measured in micrometers.

### 3.8 Bisection Assay

Twenty-five adult female *M. aquaticus* and twenty-five adult hermaphrodite *C. elegans* were placed on microscopic slides and bisected with a 25-gauge 5/8-inch hypodermic syringe needle in 40–50 µL of distilled water for *M. aquaticus* or M9 buffer for *C. elegans*. Immediately after bisection, the number of tail bends was counted over 60 seconds and at 10-, 20-, and 30-minutes post-bisection. As a control, twenty-five uncut *M. aquaticus* and twenty-five uncut *C. elegans* worms were assessed for locomotion by counting body bends.

### 3.9 Data Analysis

#### DAPI VNC neuron-like nuclei analysis

VNC neuron-like nuclei counts and densities were analyzed using GraphPad Prism 9 or R (version 4.4.2) packages dplyr, ggplot2, dunn.test, ggpubr, and openxlsx. Previously published counts for Clade 8-12 species were included in tests for VNC counts (Han et al. 2016; Stretton et al. 1978). Normality tests, including Anderson-Darling, Shapiro-Wilk, and D’Agostino & Pearson, were performed but often indicated non-normal distributions and unequal variances, largely due to small sample sizes across clades and families. Due to the lack of normality, non-parametric tests were utilized for family and clade-level analyses. The Kruskal-Wallis test was applied to assess differences in VNC neuron-like nuclei counts and densities across families and clades, while a two-sample t-test was applied to compare differences between classes using t.test() function in R. Post-hoc analyses were performed using Dunn’s test with Bonferroni correction for multiple comparisons. Families with a sample size of 1 (Aporcelaimidae, Ascarididae, Diphtherophoridae, Ethmolaimidae, and Plectidae) were excluded from family level comparison.

A simple linear regression model was fitted to examine the relationship between VNC length and VNC count using the lm() function in R (version 4.4.2) and plotted using ggplot2 package. Model assumptions were evaluated through diagnostic plots, including residuals versus fitted values to assess homoscedasticity and normal Q-Q plots to verify normality of residuals. The strength of the relationship was assessed using the coefficient of determination (R²), and the significance of the regression coefficient was evaluated using a t-test.

#### Dye-filled neuron analysis

For dye-filling assays, a two-sample t-test was applied to compare differences in total neuron counts and head neuron counts between adult and juvenile using t.test() function in R and plotted using ggplot2 package. A simple linear regression was performed to examine the relationship between body length (µm) and dye-filled neuron count for juvenile and adult separately as described above.

#### Bisection analysis

Repeated measures ANOVA was performed to compare bodybends between time using tidyverse, rstatix R packages. The time was the main factor and each subject was a repeated measures factor. Residuals were assessed for normality via the Shapiro–Wilk test and Q–Q plots.

## 4 Discussion

While we found significantly more VNC neuronal nuclei in the basal group Enoplea compared with Chromadorea, our analysis of VNC neuronal nuclei did not support the hypothesis of a one-time secondary simplification of the nervous system during the evolution of the Chromadorea. Instead, our findings suggest that neuroanatomy, as defined by number of VNC neurons, varies significantly across Enoplea (clades 1 and 2) and basal groups of the Chromodorea (clades 3-6). Furthermore, our detailed examination of *M. aquaticus* demonstrates both conserved and divergent features of nematode nervous system development and structure.

Several pieces of data prompted our original hypothesis of a reduction in the number of neurons occurring between Enoplean and Chromodorean nematodes. Our previous data and others showed that no species in Clades 9-12 had more than 76 VNC neurons (Han et al. 2016). This contrasted with the hundreds to thousands of neurons enumerated in Enoplean nematodes *Pontonema vulgare* (Malakhov 1994), *Mermis nigrescens* (Gans and Burr 1994), and *Longidorus macrosoma* (Sulston and Horvitz 1977). Indeed, in his comprehensive book on nematode structure, Malakhov (1994) suggested a general decrease in the number of neurons between Enoplida and Rhabditida. A similar suggestion was made regarding the number of sensory neurons between the paraphyletic class Adenophorea (Clades 1-6) and Secernentea (Clades 8-12) (Coomans and De Grisse 1981). However, our data do not support a one-time simplification event. Rather, we found notable differences in the number of VNC neuron-like nuclei among both the Enoplean (Clades 1-2) and basal Chromodorea (Clades 3-6) although some families showed exceptional VNC neuron counts within a class (e.g. Prismatolaimidae and Camacolaimidae). Adjusting these data for length did not modify this diversity at clade and family level comparisons. Altogether, our data suggest that rather than a one-time simplification, there appear to be multiple expansions and losses of neuron number among basal clades. The relatively more consistent number of VNC neuron-like nuclei among the more diverged Chromodorean clades (8-12) may suggest possible morphological stasis wherein this group established an evolutionary stable neuroanatomy solution that would work across habitats.

One significant limitation of our VNC data is the reliance on DAPI-stained nuclei to identify neurons, which introduces two potential sources of error. First, our VNC counts do not include possible neurons that lie within the preanal or retrovesicular ganglia. In *C. elegans,* 10 motor neurons that are logically part of the VNC are typically found within RVG (White 1976). The distribution of these neurons in other species is unknown. Second, it is possible that neuronal-like DAPI staining patterns are not representative of neurons in other species or vice versa that non-neuronal DAPI staining patterns are neurons. Lower throughput methods based on electron microscopy or immunohistochemistry may be required to confirm our findings. However, we observed variability in nuclear morphology similar to that observed in *C. elegans,* strongly suggesting our counts are generally valid.

A second limitation results from the taxa we recovered in our survey. For example, our data suggested that the Dorylaimia (Clade 2) have more neurons than the sub-class Enoplia (Clade 1). However, our samples did not include any marine species from Clade 1—work by Malakhov (1994) demonstrated that several of the Clade 1 marine species have relatively large numbers of neurons compared to our findings. We were only able to recover, stain, and identify one marine taxa, Camacolaimidae (Clade 6). The Camacolaimidae comprises a group of marine species which had notably more neurons than other basal Chromadorea taxa (Figure 3a). This may suggest that increased numbers of neurons are adaptive for marine life; however, significantly more taxa will be needed to solidify this hypothesis. Certain taxa have evidently undergone evolutionary pressures, resulting in exceptional length-to-neuron ratios. While our comprehensive survey supports the general correlation between VNC length and neuron counts among basal clades, it is important to note that this correlation does not necessarily imply causation between these two factors.

We successfully established an isogenic culture of *M. aquaticus* as a representative from the Dorylaimia (Clade 2). This culture will allow us to examine development and variation in the neuroanatomy and behavior of a basal clade species while controlling for intraspecific genetic variability. The development of the *M. aquaticus* VNC mirrored that seen in several other species wherein a large number of neurons are added to the VNC during the early stages of post-embryonic development with no subsequent additions (Han et al. 2016; Sulston 1976). This contrasts with anatomical data from Enoplida (Clade 1), suggesting a lack of constancy in neuron numbers (Malakhov 1994). Interestingly, the dye-filling data suggested contrasting patterns of development. While many of the anterior neurons dye-filled throughout *M. aquaticus* post-embyronic development, the body wall dye-filled neurons increased in number along with body size.

The *M. aquaticus* body wall neurons shown in our dye-filling data are distinct from previous dye-filling results in Chromadorea (Han et al. 2016; Hong et al. 2019; Tong and Bürglin 2010). Determining homologs of these neurons in *C. elegans* neurons will be challenging and likely require additional molecular characterization. The most similar neuron class in *C. elegans* are the two pairs of anterior and posterior deirid neurons. While the deirid neurons of *C. elegans* have ciliated endings, these are not exposed to the environment and do not dye-fill in wild-type animals. Unlike *C. elegans,* ciliated body wall neurons have been reported from several species in other basal clades (Hope et al. 1982; Malakhov 1994; Wright and Carter 1980). An ultrastructural examination of the body wall neurons in *M. aquaticus* could provide insight into their homology.

Our finding that bisected *M. aquaticus* tails are able to continue movement long after they have been bisected from connection to the central nerve ring may suggest a more distributed neuronal network. An alternative explanation for our bisection data is that *M. aquaticus* is better able to hold individual cells in place following a traumatic event. While *M. aquaticus* has a relatively large number of neurons, we didn’t observe any obviously unique movements in intact animals. Although *M. aquaticus* is a predator, unlike the bacterial feeding *C. elegans,* we do not think this would require the larger number of neurons inferred from our data. Indeed, *P. pacificus,* which is a much closer relative to *C. elegans,* also can feed as a predator, but does not display a corresponding increase in number of neurons (Cook et al. 2025; Han et al. 2016).

The *C. elegans* nervous system is considered a small-world network (Watts and Strogatz 1998). Perhaps the decentralized control seen here reflects an ancestral condition that persists in basal nematodes which offers adaptive advantages in fluctuating environments. However, this currently speculative hypothesis will require additional connectivity data.

## 5 Conclusions

Our comprehensive evaluation of VNC neuroanatomy across diverse nematode taxa revealed substantial variation in neuron numbers with greater neuroanatomical diversity within nematodes than previously recognized. Contrary to the hypothesis of a one-time evolutionary simplification, our results indicate complex evolutionary trends in neuron numbers across different taxa. In general, basal Enoplean nematodes showed significantly greater VNC neuron counts than the derived taxa such as *C. elegans.* However, our established isogenic line of *M. aquaticus* showed similar patterns of neuronal development in the early-stage juvenile like derived species. Notably, *M. aquaticus* also demonstrated extensive sensory neuron distribution in the head and along its body length, which is distinct from the restricted dye-filling of head sensory neurons observed in diverged species. Bisection assays using *M. aquaticus* further suggests that basal nematodes may possess less centralized nervous system.

## Acknowledgments

We would like to acknowledge Dr. Holly Bik and Dr. Tiago José Pereira for providing marine nematode samples. ML was supported by a Jonathan Baldwin Turner fellowship. The Schroeder lab acknowledges support from NIH (OD010943) and USDA-NIFA (2021-67013-33737).

